# An amended potassium persulfate ABTS antioxidant assay used for medicinal plant extracts revealed variable antioxidant capacity based upon plant extraction process

**DOI:** 10.1101/2020.07.15.204065

**Authors:** Cody E. Mingle, Anthony L. Newsome

## Abstract

Use of potassium persulfate (K_2_S_2_0_8_) for oxidation of 7.0 mM ABTS to a stable ABTS radical for antioxidant studies was first reported in 1999. A feature of this popular antioxidant assay has been the requirement of an overnight reaction (6 to 12 h) for the formation of a stable ABTS colored radical. It is now reported that when the concentration of ABTS is lowered to 0.7 mM, complete oxidation to the stable cation radical occurs in 30 min, thus circumventing the necessary overnight step. Using this format, it is now possible to accurately assess antioxidant activity based on the potassium persulfate/ABTS format in less than one hour which includes formation time of a stable ABTS radical. This methodology documented the presence of antioxidant properties of plant extracts used in Traditional Chinese Medicine. The degree of antioxidant activity was directly related to the extraction method. Greater antioxidant activity was associated with butanol extraction. When incorporated into a microtiter plate format, it supported rapid assessment of multiple determinations of dilutions of plant extracts in less than one hour which included time required for formation of a stable ABTS radical. The ease, improved time prerequisites, and minimal reagent needs with the microtiter plate format, makes this design attractive. It would prove of particular interest to individuals engaged in both smaller and high-volume throughput antioxidant assays of food and health products, and other biological fluids and tissues.

## Introduction

Assessment of antioxidant activity in foods, health products, natural products used for their purported benefit to humans, and other biological fluids and tissues continues to be an active pursuit in the scientific community. The ABTS assay has gained acceptance as a valuable tool in the assessment of antioxidant activity and pursuit of these studies. The ABTS colored cation radical is generated upon the loss of an electron by the nitrogen atom of ABTS [2,2’-azino-bis(3-ethylbenzthiazoline-6-sulphonic acid)] [1]. For antioxidant assay purposes, this reaction (oxidation and loss of an electron and formation of the ABTS cation radical) can be brought about by use of potassium persulfate (K_2_S_2_O_8_) [2]. With use of potassium persulfate, the ABTS radical was generated directly with no intermediary radical. Thus, the ABTS radical is preformed prior to addition of test antioxidants, eliminating the possibility that test substances could interfere with generation of the ABTS radical [2]. Use of potassium persulfate for generation of ABTS radical has garnered popularity [2-5]. Spectrophotometer readings at absorbance setting of 734 nm were regularly used and Trolox has often served as an antioxidant standard [2-5].

A feature of this assay system has been the use of 2.45 mM potassium persulfate in conjunction with 7.0 mM of ABTS for ABTS radical generation. This has been accompanied by overnight (12-16 h) incubation and subsequent dilution of the ABTS radical is to a working absorbance value of approximately 0.7 (or slighter higher in some studies) at 734 nm [2-5]. Working assay volumes were typically 1.0 ml (or greater in some studies). Absorbance and antioxidant activity are most often determined within the first 6 min after addition of antioxidant [2-4].

Described in this current study is the use of 0.7 mM ABTS (rather than the previously described use of 7.0 mM ABTS) in conjunction with 2.45 mM potassium persulfate. When this scheme was followed, the generation of the ABTS radical went to completion in 30 min, thus avoiding the previously described overnight (12-16 h) incubation period. The resulting ABTS radical was stable and amenable to incorporation to a microtiter plate format.

This newly described method afforded the opportunity to initiate and assess antioxidant activity in less than one hour, which included generation time for radical formation and assessment of test antioxidants. The microtiter plate format afforded the ability to more easily perform serial dilutions and multiple determinations (triplicate) of test compounds. An additional benefit of this modified assay is the reduced amount of test reagents needed to access antioxidant activity as compared to the use of ml test volumes previously described [2-5].

This format served as a practical means to rapidly assess antioxidant activity in extracts of plants that are popular diet supplements consumed in the form of tea, especially in China. Test results demonstrated antioxidant activity and variability among the different plant extracts. Multiple determinations of antioxidant activity of serial dilutions of plant extracts based on this format were consistent and reproducible. This modified and more rapid ABTS colorimetric assay for determination of antioxidant activity also has potential broad applications for assessment of antioxidant activity in a variety of other biological specimens.

## Materials and methods

### ABTS radical preparation protocol

ABTS (2,2′-azino-bis(3-ethylbenzothiazoline-6-sulfonic acid) was dissolved in 1X phosphate buffered saline (HyClone Laboratories, Logan UT). For this, 5mg ABTS (Roche) tablets (Sigma-Aldrich, St. Louis, MO) were added to 7 ml PBS which yielded a final concentration of 0.7 mM ABTS. A 2.45 mM solution of potassium persulfate (Sigma-Aldrich, St. Louis, MO) was prepared in deionized water. A volume of 7.0 ml of ABTS was added to 7.0 ml of potassium persulfate in a glass tube and placed in the dark at room temp (22 °C). The starting ABTS solution was prepared fresh before use and the potassium persulfate prepared fresh every 2 weeks. Antioxidant assessment was performed immediately after oxidation of ABTS with complete formation of the ABTS radical.

### Microtiter plate preparation

While the ABTS radical formation was in progress (30 min), test microtiter (96 well polystyrene clear flat bottom) plates (Denville Scientific Inc., Charlotte, NC) were prepared. For this 50 µl of PBS was added to all wells except for the outside row of wells on each side of the microtiter plate. A volume of 50 µl of test compound was added to the first test well (total volume 100 µl). Fifty µl was removed and serially diluted across the microtiter plate row. The final 50 µl from the last test well was discarded. A multichannel pipette (VistaLab Technologies, Brewster, NY) was used to perform tests in duplicate, triplicate, or whatever number of multiple repeats were deemed appropriate.

### Test compounds

Plant extracts were provided by Guangxi Botanical Garden of Medicinal Plants (Nanning, PR China). The solvents petroleum ether, ethyl acetate, butanol, or chloroform were used for preparation of extracts. Preparations were taken to dryness and subsequently suspended in dimethyl sulfoxide (DMSO) at a concentration of 10 mg/ml. DMSO only served as a negative control for the antioxidant studies. Trolox (6-hydroxy-2,5,7,8-tetramethylchroman-2-carboxylic acid) (Sigma-Aldrich, St. Louis, MO) served as an antioxidant standard and it was likewise suspended in DMSO at 10 mg/ml.

### Test assay

Once dilutions of test compounds were prepared on the microtiter plates, 150 µl of the ABTS radical was added to each well (total volume 200 µl/well) with a multichannel pipette to give an initial absorbance of 0.7 at 734 nm (CLARIOstar Microplate Reader, BMG Labtech Inc., Cary, NC). Absorbance readings (at 734 nm) were determined at room temp at 5, 15, and 30 min after addition of the ABTS radical to test compound dilutions. Antioxidant activity of test compounds, Trolox, and DMSO controls were expressed as percent inhibition of absorbance (Ia), using the following equation:

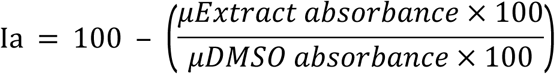

## Results

Incorporation of 0.7 mM (rather than the previously described use of 7.0 mM) ABTS resulted in oxidation to completion of ABTS to the ABTS cation radical in 30 min as noted by absorbance at 734 nm (Fig.1). The newly formed radical was suitably stable for the assessment of antioxidant activity of test compounds. Assessment was performed immediately (within 30 min) after oxidation of ABTS to the ABTS radical.

**Fig 1.**
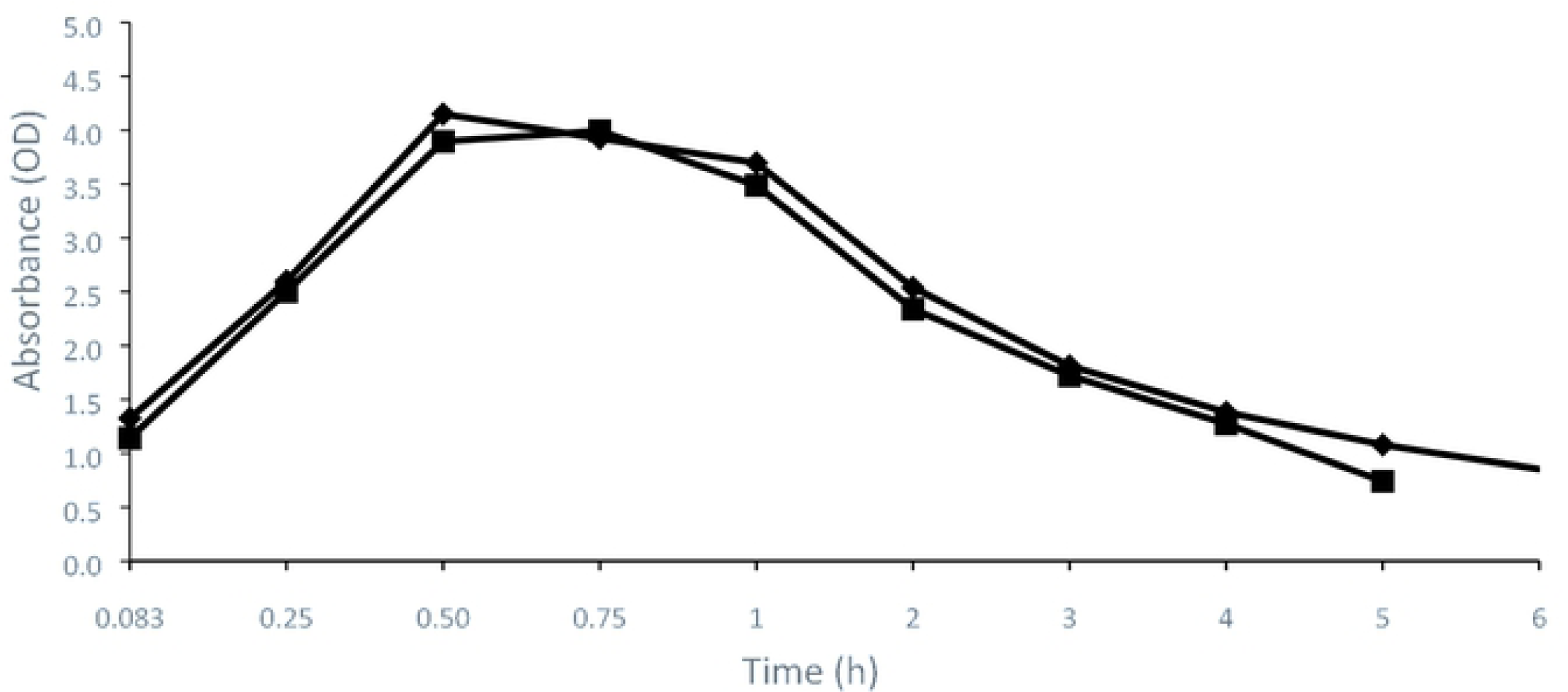
Conversion of 0.7mM ABTS to ABTS^+^ with 2.45 mM potassium persulfate. Values represent average of triplicate determinations. The SD was consistently less than 0.11. Deionized Water (▪), PBS (♦).

Antioxidant activity (percent inhibition of absorbance of ABTS radical) of plant extracts and the Trolox standard was apparent after 5 min. (Figs. 2-4). Additional measurements of antioxidant activity of up to 30 min did not yield substantial differences from that observed at 5 min (data not shown). Thus, figures represent antioxidant activity observed at the 5 min time point. Antioxidant activity of plant extracts varied among the plants tested and the plant extraction methods. The DMSO only microtiter test wells displayed negligible antioxidant activity. Antioxidant activity of extracts from *Cyclocarya paliurus* was apparent after 5 min (Fig. 2).

**Fig 2.**
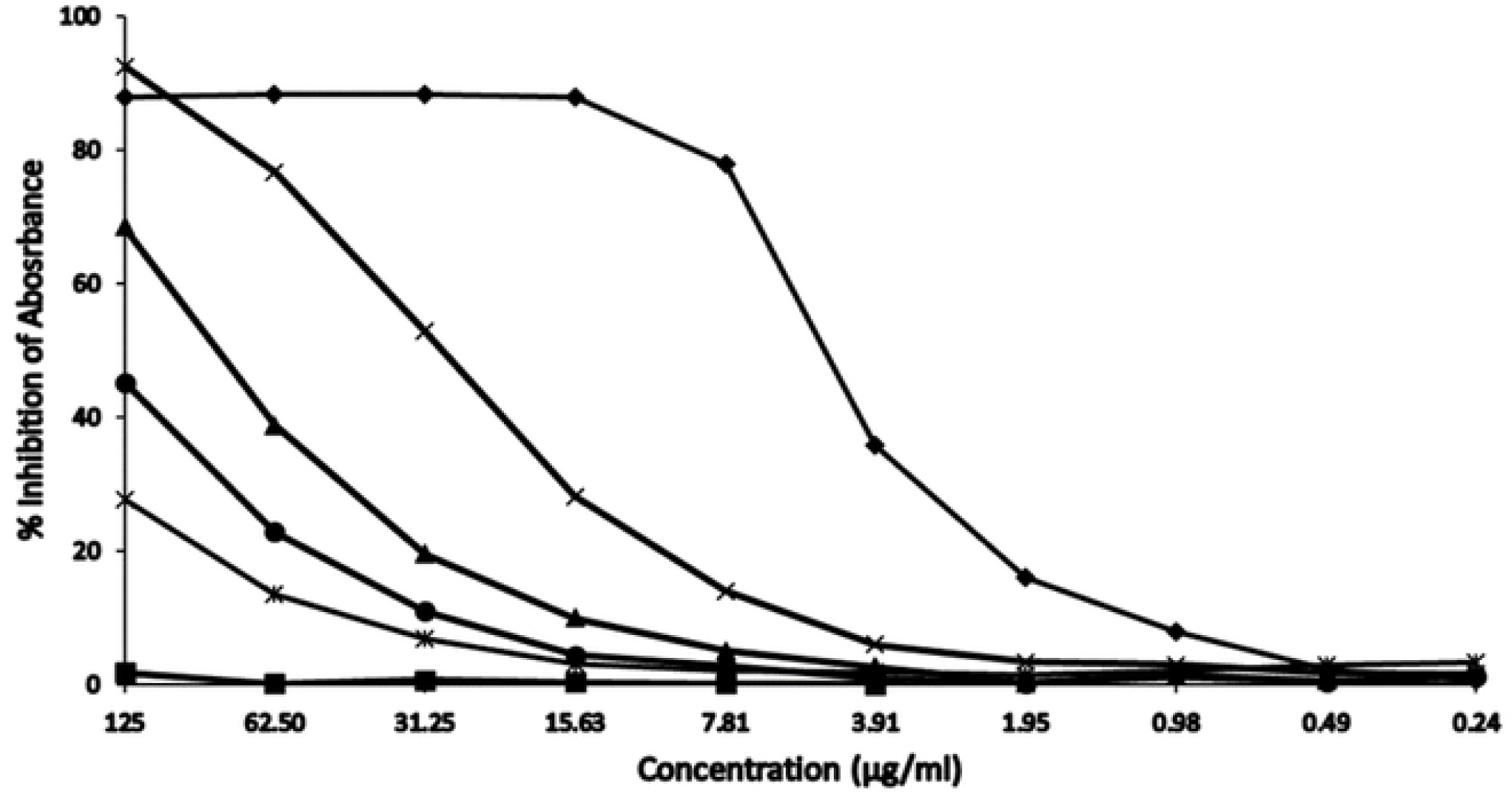
*Cyclocarya paliurus* extract inhibition of absorbance on ABTS cation radical. Values represent average of triplicate determinations. The SD was consistently less than 0.084. Petroleum ether (*), Ethyl acetate (▲), Butanol (X), Chloroform (•), Trolox (♦), DMSO (▪).

All extracts displayed antioxidant activity at the highest concentration tested (125 µg/ml). The butanol extract displayed the greatest antioxidant activity and was equivalent to that of Trolox at the highest concentration (125 µg/ml). Its antioxidant activity remained greater than that of the other extracts (petroleum ether, ethyl acetate, and chloroform) per dilution to lower concentrations on the microtiter plate. Following a 2-fold dilution to 62.50 µg/ml in the microtiter plate well only the butanol extract displayed greater than 40 percent inhibition of absorbance. The Trolox standard displayed a 40 percent or greater inhibition of absorbance to a concentration of approximately 3.91 µg/ml. Antioxidant activity of extracts of *Gynostemma pentaphyllum* was likewise demonstrated (Fig. 3).

**Fig 3.**
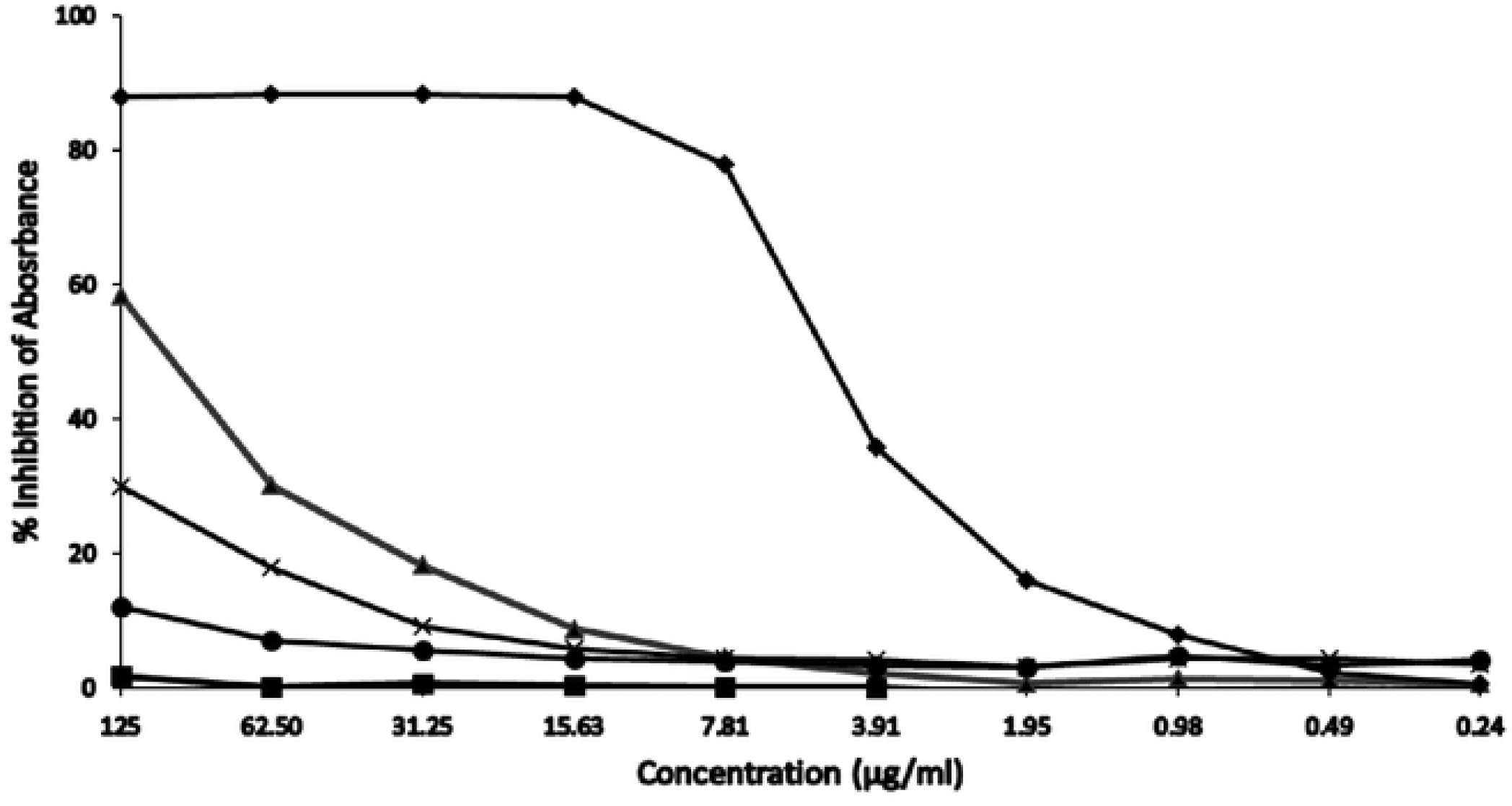
*Gynostemma* pentaphyllum extract inhibition of absorbance on ABTS cation radical. Values represent average of triplicate determinations. The SD was consistently less than 0.092. Ethyl acetate (▲), Butanol (**X**), Chloroform (•),Trolox (♦), DMSO (▪).

Greatest antioxidant activity was associated with the ethyl acetate extract. Even at the highest concentration tested (125 µg/ml), inhibition of absorbance did not exceed 60 percent. Less than 40 percent inhibition was observed in all the *G. pentaphyllum* extracts at 62.50 µg/ml or less (Fig. 3). A characteristic of the antioxidant activity of extracts from *Usnea diffracta* was that the greatest antioxidant activity was again associated with the butanol extract (Fig. 4).

**Fig 4.**
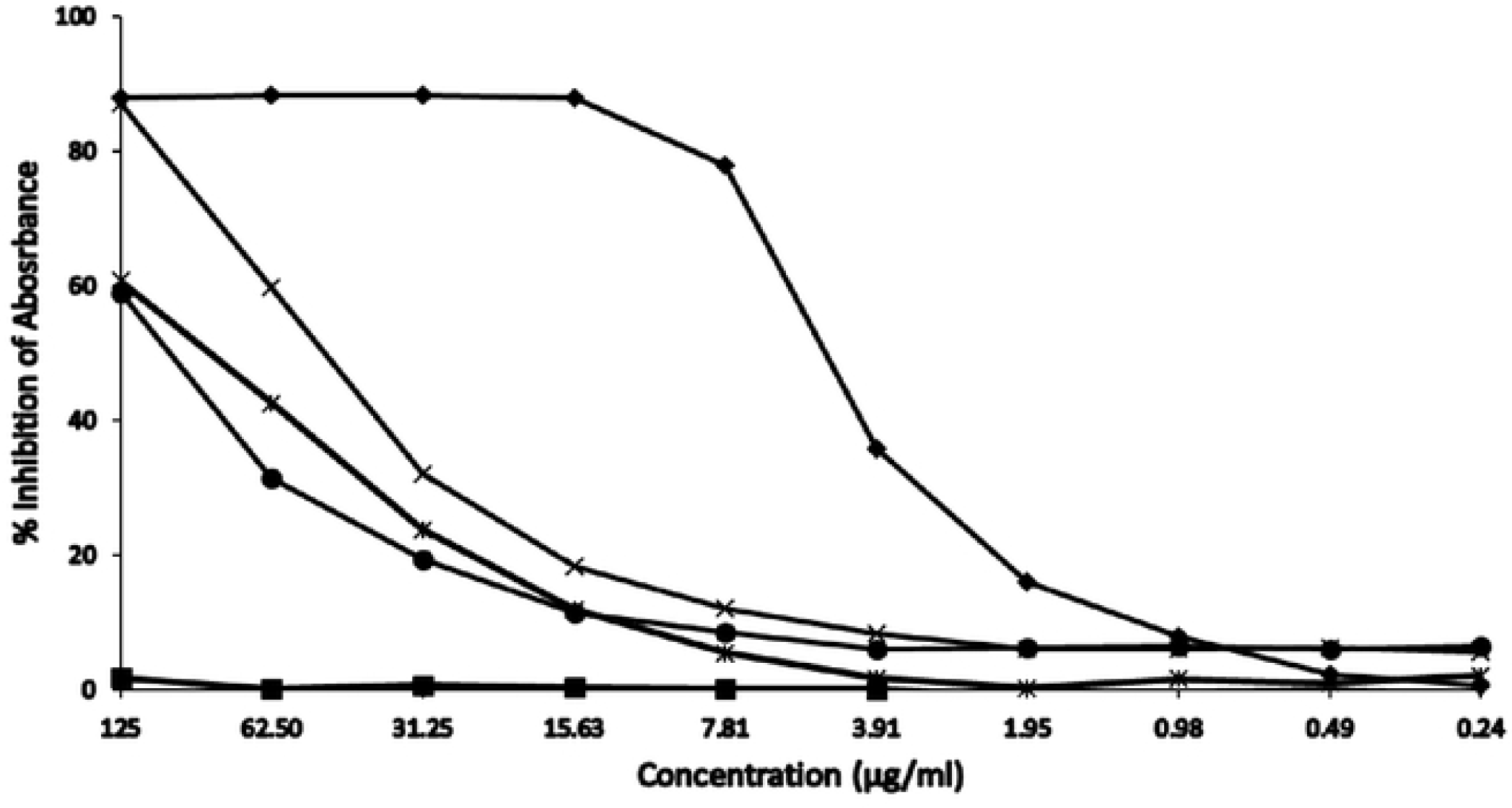
*Usnea diffracta* extract inhibition of absorbance on ABTS cation radical. Values represent average of triplicate determinations. The SD was consistently less than 0.018. Petroleum ether (*), Butanol (**X**), Chloroform (•), Trolox (♦), DMSO (▪).

At its highest concentration (125 µg/ml) it was equivalent to that of Trolox. Following a 2 fold dilution to 62.50 µg/ml, greater than 40 percent inhibition of absorbance was still apparent in the butanol and chloroform extracts.

A feature of the antioxidant activity of all the plant extracts compared to Trolox was the rapid diminishment of antioxidant activity of plant extracts following 2-fold dilutions in the microtiter plate wells. Trolox antioxidant activity continued to be apparent following the 2-fold serial dilutions and a nearly 40 percent inhibition of absorbance was displayed by Trolox at a concentration of 3.91 µg/ml. Antioxidant activity was not associated with DMSO alone control microtiter plate wells.

## Discussion

The newly described format functioned effectively for assessment of antioxidant activity of plant extracts that are used for human benefit. This includes treatments of a variety of diseases and previous studies have shown the plant extracts to have a variety of pharmacologic properties. Studies have also shown some of the plant used in this study can exhibit antioxidant properties.

The primary application *Cylocarya paliurus* has been use of its leaves in tea production and antioxidant properties have been ascribed to water soluble extracts of the leaves. Among 21 different geographic populations of the plant, there was a significant difference in antioxidant activities [9]. An ethanol extract of its leaves has exhibited cytotoxicity against human lung and breast cancer cells [10]. Chinese medicine and treatment of a variety of diseases including cancer has been ascribed to Gynostemma pentaphyllum (Thunb.) Makino. It has displayed different pharmacological properties that include antioxidant properties [11]. The usnic acid and diffractic acid from the lichen *U. diffracta* have described analgesic and antipyretic components [12]. Antioxidant activity has been demonstrated in other species within this genus [13].

In the current study, the greatest level of antioxidant activity in two of the three plant extracts was associated with extraction by butanol. At the highest concentration tested (125 µg/ml) antioxidant activity was comparable to that of a Trolox standard (also at 125 µg/ml). The antioxidant activity of the plant extracts rapidly diminished upon dilution in the microtiter plate format. The Trolox standard, however, maintained prominent antioxidant activity upon dilution to much lower concentrations (µg/ml).

Generation of the ABTS radical using potassium persulfate is a conventional approach for assessment of antioxidant activity. The ABTS and potassium persulfate react in a stoichiometric manner yielding the ABTS cation radical. Previously, this approach has carried with it a requirement to allow at least 6 h, and typically a 6 to 12 h overnight reaction for oxidation completion and formation of the stable ABTS radical cation for antioxidant studies [2-5].

In the current study, the capability to generate a suitably stable ABTS cation radical using potassium persulfate in 30 minutes for antioxidant studies provides a logistic alternative to the previously described 6 h minimum reaction time [2-5]. Furthermore, the ABTS radical was readily adaptable into a microtiter-based format providing an additional advantage of easily performed repetitive testing of different dilutions of test antioxidant.

Assessment of antioxidant activity in body fluids, nutritional products, and chemical systems continues to be an important scientific activity. With the continued scientific interest in assessment of antioxidant activities, there also continues to be efforts directed at efficient and cost effective protocols for assessment of antioxidant activities and high throughput microtiter based assays have been incorporated into antioxidant assays [6-8].

## Conclusions

Outcomes of this study showed that extraction methodology provided an important role in assessment of antioxidant capacity of plant extracts. This study was accompanied by incorporation of a much more rapid (1 h) alternative for determinations of antioxidant activity using the ABTS/potassium persulfate-based format. The incorporation of a microtiter-based format additionally supports rapid assessment of multiple determinations of test dilutions of antioxidant compounds. In addition to use in dietary plant extracts as described in this study, it also provides potential for use in a variety of chemical and biological compounds such as foods and health products This study may also have broad potential for use in clinical applications such as determination of antioxidant activity in serum, urine, and stool extracts.

## Acknowledgements

Appreciation is expressed to Dr. Iris Gao and Guangxi Botanical Garden of Medicinal Plants (GBGMP), Nanning, Guangxi, China, for preparation of the plant extracts used in the study.

